# Free energy calculations suggest a mechanism for Na+/K+-ATPase ion selectivity

**DOI:** 10.1101/106724

**Authors:** Asghar M. Razavi, Lucie Delemotte, Joshua R. Berlin, Vincenzo Carnevale, Vincent A. Voelz

## Abstract

Na+/K+-ATPase transports Na^+^ and K^+^ ions across the cell membrane via an ion binding site made alternatively accessible to the intra- and extracellular milieu by conformational transitions that confer marked changes in ion binding stoichiometry and selectivity. To probe the mechanism of these changes, we used molecular simulation approaches to identify the protonation state of Na^+^ and K^+^ coordinating residues in E1P and E2P conformations. Further analysis of these simulations revealed a novel molecular mechanism responsible for the change in protonation state: the conformation-dependent binding of an anion (a chloride ion in our simulations) to a previously unrecognized cytoplasmic site in the loop between transmembrane helices 8 and 9, which influences the electrostatic potential of the crucial Na^+^-coordinating residue D926. This mechanistic model is consistent with experimental observations and provides a molecular-level picture of how E1P to E2P enzyme conformational transitions are coupled to changes in ion binding stoichiometry and selectivity.

---

Na+/K+-ATPase is a membrane protein that actively transports sodium ions out of the cell while importing potassium ions, both against their electrochemical gradients, thus providing potential energy necessary for many essential cellular functions (1-3). Malfunction of Na+/K+-ATPase has been linked to numerous diseases including impaired memory and learning, familial hemiplegic migraine 2, rapid-onset dystonia Parkinsonism, and heart failure (4-6), thus, Na+/K+-ATPase is an important target for treatment of brain and heart conditions (7,8).

By harnessing chemical energy stored in ATP, the Na+/K+-ATPase cycles between two major conformational states during active ion pumping: one state with high affinity for sodium ions (E1), and a second with high affinity for potassium ion (E2). While the complete mechanism of function of the Na+/K+-ATPase is more complicated, several conformational transitions described more fully in involving several conformational transitions described more fully in the Post-Albers scheme (9-11), here we focus on the sodium-bond E1P and potassium-bound E2P states, for which crystal structures are available (12-17). One of the central features of ion transport by the Na+/K+-ATPase is a change in ion selectivity between E1 and E2 conformations (3). The origin of this selectivity change has been a focus of Na+/K+-ATPase research since its discovery by Jens Christian Skou in 1957 (1). Extensive mutational studies have identified key residues involved in ion binding, which include five acidic residues: Asp804, Asp808, and Asp926, Glu327 and Glu779 (13) whose orientation, distance, and ion coordination have been proposed to be involved in determining selectivity (3). The first crystal structure of the Na+/K+-ATPase, solved for the potassium-bound E2P state in 2007 (15) and later followed by several higher resolution structures (14,16,17), revealed a binding site in which these five acidic residues and several backbone carbonyl oxygens tightly coordinate two potassium ions within a ~15 Å^3^ space. The close proximity of acidic binding residues (Figure 1) suggested that some of these residues must be protonated. Indeed, protonation of binding site residues is intrinsic to the transport cycle of the Ca^2+^-pump, SERCA, a closely related P-Type ATPase (18).

**FIGURE 1.**
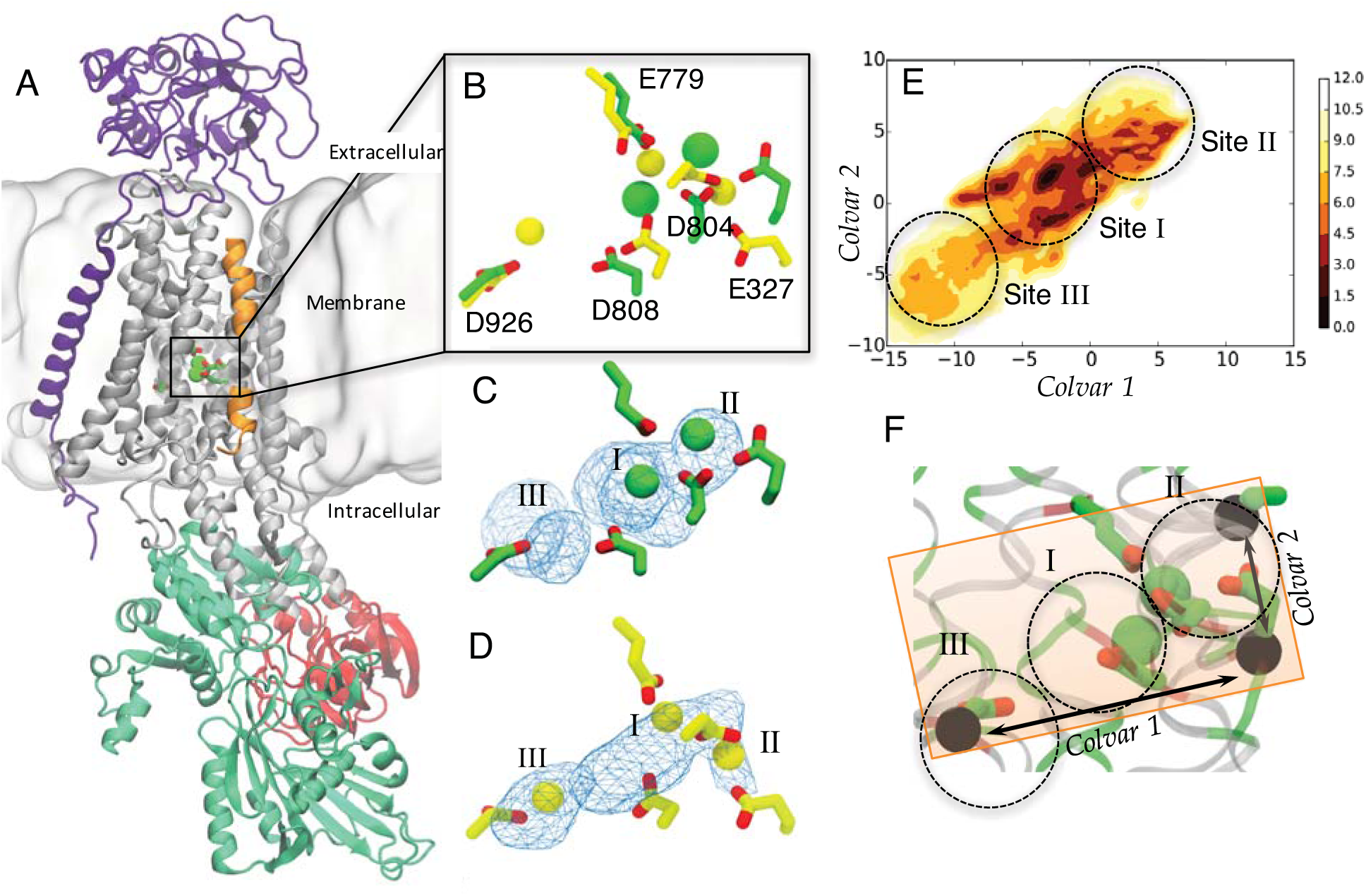
Molecular structure of Na+/K+-ATPase. (A) The α-subunit is shown in gray, with its cytoplasmic domains colored: A-domain (red), N-domain (blue) and P-domain (green). The β-subunit is shown in violet, and γ-subunit shown in orange. (B) A superposition of the binding site for sodium-bound E1P (yellow) and potassium-bound E2P (green). (C and D) Density of ions sampled in 200 ns simulations (blue mesh) superimposed on crystal structures for potassium (C) and sodium (D). (E) A 2D potential of mean force (PMF) calculated from metadynamics simulations, for one potassium ion in the E2P binding site when D926 is protonated. Approximate location of sites I, II, and III are highlighted with circles. (F) Scheme representing the collective variables used to calculate the PMF (see Experimental procedures). Black circles show the reference atoms (alpha carbons) used for the metadynamics simulation.

The notion that protonation states can modulate ion selectivity is strongly supported by a joint computational and experimental study by Yu et al. in which different protonation states of potassium-bound E2P state were predicted to confer differences in ion selectivity (19). In this work, electrophysiological experiments confirmed computational predictions that potassium selectivity decreases with increasing pH, consistent with previous pH studies (20). However, the exact protonation state could not be determined, nor could the results provide insight into the protonation state of sodium-bound conformations. Most importantly, it is not clear how changes in protonation state are controlled by or coupled to structural changes during active transport of ions. The first crystal structures of the sodium-bound E1P state were obtained in 2013 (12,13), providing an unprecedented opportunity to discover the molecular origins of selectivity in atomic detail – now a half-century-old question.

Here, we address the question of how ion selectivity and binding stoichiometry are achieved at the ion binding sites in different protein conformations by extensively scrutinizing Na+/K+-ATPase in atomic detail using molecular simulation. Our results suggest that the protonation state is different in the sodium- and potassium-bound states. We show that while the protonation state controls potassium ion selectivity and stoichiometry, selectivity for sodium appears to be determined by the steric constraints imposed by the binding site geometry and independent from protonation state. Our simulations also suggest the presence of a previously unknown cytoplasmic binding site for anions that helps control the change in protonation state of Asp926, a crucial residue in forming the sodium-specific ion binding site III. Furthermore, we show that access to this cytoplasmic anion binding site is largely controlled by conformational changes in the cytoplasmic loops between TM6-TM7 and TM8-TM9, which is different in sodium-bound E1P versus potassium-bound E2P states.

## Results and Discussion

The sidechain carboxylate oxygens of five acidic amino acids (E327, E779, D804, D808, and D926) are critical in coordinating Na^+^ and K^+^ in the Na+/K+-ATPase. D926 is located on the transmembrane (TM) helix 8 (TM8), and E327 in TM4 (12), while the other acidic residues are located on TM5 and TM6 and are considerably closer to each other (Figure 1). These five acidic residues form three binding sites: sites I and II are formed by E327, E779, D804 and D808, along with backbone carbonyl moieties of TM4 residues, and can be occupied by either sodium or potassium ions; site III is formed by D926, D808 and oxygen atoms of TM5 residues, and is only occupied in the E1P state by Na^+^ (12,13). As in previous computational studies of Na+/K+-ATPase (19), we first tried to predict the protonation state of titratable residues using the PROPKA algorithm, an empirical method which relates desolvation effects and intra-protein interactions to the positions and chemical nature of residues proximate to the titratable sites (21). According to these calculations, only D804 and E779 are protonated in E1P, but for the E2P state all are protonated except D808. (Figure S1). However, our molecular dynamics simulations initialized from these configurations produced, after 100 ns of equilibration, configurations inconsistent with these predictions: self-consistency tests applying PROPKA to the snapshots extracted from these equilibrated trajectories predicted protonation states significantly different from those in the initial structure (Figure S1). Unsatisfied by these inconsistent results, we turned to more extensive, yet more accurate, simulation-based methods to determine the protonation states for both sodium-bound E1P and potassium-bound E2P states.

### Site III has a high affinity for both sodium and potassium ions when D926 is deprotonated

With five acidic binding residues, systematic comparison of the 2^5^ = 32 possible protonation combinations for both the sodium-bound E1P and potassium-bound E2P states of Na+/K+-ATPase is a challenging task, requiring molecular dynamics simulations for a quarter of a million atoms for each possible protonation state. Fortunately, several protonation states could be eliminated. Our initial simulations of 210 ns, in which none of the acidic binding residues were protonated, displayed significant ion-binding conformational inconsistency from crystal structures. In these simulations, both sodium and potassium ions relocated preferentially to site III (Figure 1C and 1D), while all crystal structures of potassium-bound states show that only sites I and II are occupied by potassium ions (14-17). Since the carboxylate moiety of D926 interacts exclusively with the cation bound to site III; we hypothesized that the protonation of this residue in the potassium-bound E2P state might result in detachment of the potassium ion from site III. To test this hypothesis, we continued the simulation after protonation of D926. Within several nanoseconds, the potassium ion bound in site III migrated towards sites I and II. We furthermore used metadynamics simulations (see Experimental procedures) to calculate potential of mean force (PMF) along two collective variables (22) characterizing the bound ion density; the results revealed that protonation of D926 leads to a more favorable binding of K^+^ to binding sites I and II and not to site III (Figure 1E). Occupation of site III by Na^+^ ions, on the other hand, is consistent with the presence of such a cation in the crystal structure, thus leading us to conclude that D926 should be deprotonated in the Na^+^-bound E1P state, but not in the K^+^-bound E2P state. A similar conclusion was reached by Nissen and co-workers (5).

### The sodium-bound E1P and potassium-bound E2P conformations have distinct protonation states

Next, we evaluated the remaining 2^4^ = 16 protonation state possibilities for the acidic binding residues: E327, E779, D804, and D808. As a first step, we tested the possibility that all four residues are deprotonated or protonated in both the E1P and E2P conformations. Not surprisingly, in equilibration simulations of each case, the large charge density in the binding site resulted in the binding of counter-ions from the solution buffer to neutralize the binding site charge, leading us to reject both possibilities (data not shown). For the remaining 14 combinations (Table S1), we were able to filter the results of equilibrium MD simulations using structural criteria, namely the RMSD of binding-site ions and residues from crystal structures, the degree of fluctuation of ions in the binding site, and ion coordination numbers (Figures S2 and S3). This comprehensive analysis resulted in the elimination of seven additional combinations in the sodium-bound E1P state and eight additional protonation combinations in the potassium-bound E2P state (Tables S2 and S3).

For the remaining six and seven protonation combinations in the potassium- and sodium-bound conformations, respectively, we used free energy perturbation (FEP) methods to determine the relative free energy of each protonation state (see Experimental procedures). To perform these calculations efficiently, alchemical intermediates were constructed along a network of pathways bridging each protonation state in the potassium- and sodium-bound conformations (Figure 2). Forward and backward transformations, each starting from independent equilibrated structures to avoid bias, were used to validate the accuracy of these calculations. In total, this amounted to ten FEP calculations for the potassium-bound E2P state (five forward and five backward), and twelve FEP calculations for the sodium-bound E1P state (six forward and six backward). Moreover, in simulations for which the total charge of the system changes during FEP, a reference state is used to compensate for anisotropic electrostatic effects (see Experimental procedures). Predicted free energy differences, Δ*G*, for all FEP simulations are summarized in Figure 2. The results show that only two residues out of four (D804, D808, E327, E779) need to be protonated in the sodium-bound and potassium-bound conformations, although the preferred protonated residues are different in each case. Specifically, for the sodium-bound E1P state residues E779 and D804 are most likely to be protonated, while for potassium-bound E2P state residues E327 and D808 are most likely to be protonated (Figure 2).

**FIGURE 2.**
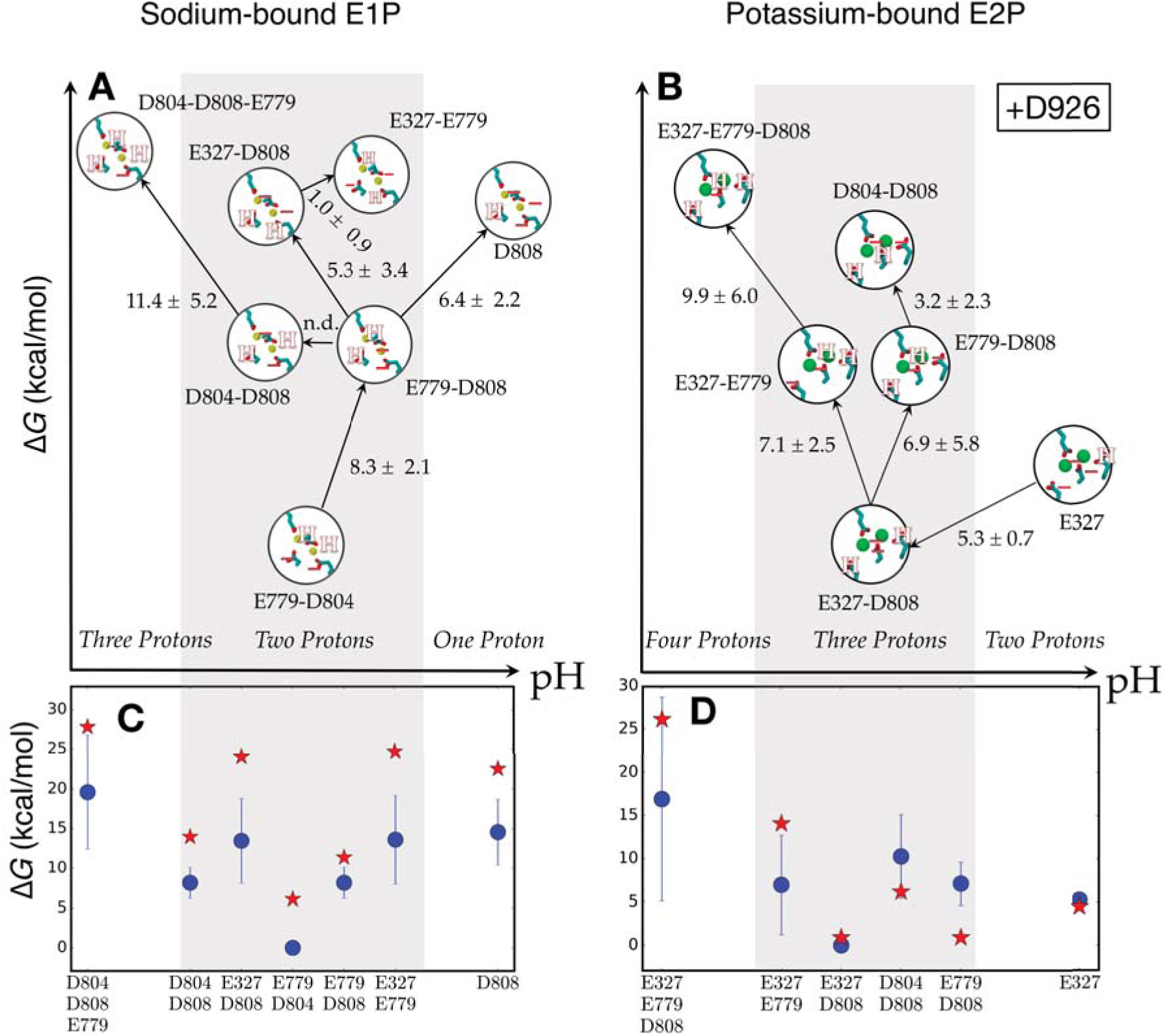
FEP calculations of relative free energies of binding-site protonation states for the sodium-bound E1P (A) and potassium-bound E2P (B) conformations of the Na+/K+-ATPase. Protonated residues are labeled for each state, and indicated by a letter ‘H’ in each cartoon. n.d.: not determined. The inset in panel B is a reminder that D926 is protonated in all these protonation states (see text). One, two, three, or four protons denote that 1, 2, 3, or 4 acidic residues in the binding site are protonated, respectively. (C and D) Blue circles show the relative free energy of each protonation state obtained from corresponding top panels, with error bars. Red stars show, for each protonation state, the change in free energy incurred about transforming (C) three sodium ions into three potassium ions, and (D) two potassium ions to two sodium ions.

### FEP-determined protonation states correctly reproduce experimentally measured ion binding affinities

We performed further alchemical FEP calculations (see Experimental procedures) to determine the relative affinities of sodium and potassium ions for the binding site in E1P and E2P conformations, for the most likely protonation states described in the previous section (Figure 2). In these calculations, sodium ions bound in the E1P state are alchemically transformed into potassium ions; likewise, potassium ions bound in the E2P state are transformed into sodium ions (see Experimental procedures). The results correctly predict the ion-binding selectivity of E1P and E2P states, estimating a preference of approximately 6 kcal/mol for sodium over potassium ions in the E1P conformation, and a preference of approximately 1 kcal/mol for potassium over sodium ions in the E2P conformation. The experimentally measured selectivities are 1.5 kcal/mol preference for sodium in E1P, and 4.5 kcal/mol preference for potassium ions in E2P (3,23,24). While these experimental measurements are apparent affinities and cannot be directly compared to our calculated affinities, nevertheless, it is noteworthy that the calculation of ion affinities by FEP qualitatively reproduces ion selectivity in both E1P and E2P states.

### Molecular mechanism of selectivity in Na+/K+-ATPase

While the results above provide a description of distinct differences in protonation states and ion binding selectivity for the E1P versus E2P conformations, a more important question remains: What are the structural mechanisms underlying these distinct differences between the electrostatic environments of E1P and E2P? To provide insights, we used alchemical free energy calculations to determine the affinity of Na^+^ versus K^+^ in the E1P and E2P binding sites, as well as electrostatic decomposition analysis to discover the origins of pKa shift for the binding site residues in E1P and E2P states.

#### Selectivity for sodium ions in the E1P state arises from binding site volume constraints

For all possible protonation states that show low deviations from crystal structures, i.e. the seven protonation states for E1P state and 6 protonation states in E2P state shown in Figure 2A and 2B, we calculated the binding free energy of sodium versus potassium to determine which ions are more preferred in each of these protonation states (Figure 2C and 2D). Our FEP results predict that in the E1P conformation the affinity for sodium ions relative to potassium ions does not depend on the specific protonation state. Regardless of protonation state, the E1P state always has higher affinity for sodium than potassium ions (Figure 2C and Figure S4). Seemingly, only two reasons can justify this systematic increase in free energy upon change of sodium into potassium: a lack of ligands (water or protein oxygens) leading to an incomplete shell of solvation for the potassium ion, or unfavorable repulsive van der Waals interactions between the binding pocket and the bigger (compared to sodium) solvated ion shell of potassium.

We found that the total coordination number increases as sodium ions are alchemically transformed to potassium ions, reaching values typical of a potassium ion in solution (Figure S5), therefore, it is unlikely that potassium ions are suffering from lack of coordinating ligands. On the other hand, the number of coordinating water molecules also increases as sodium ions are transformed into potassium ions (Figure S5). These observations suggest that the E1P binding site cannot accommodate three larger solvated potassium ions. Accordingly, by analyzing separately each contribution to the average potential energy in two distinct simulations of the sodium- and potassium-bound states, we concluded that Lennard Jones interactions have a destabilizing role (by as much as 8 kcal/mol) in passing from the first to the second case. We note that the same energy difference for an ion in aqueous environment has the opposite sign (it is approximately -2.5 kcal/mol, see table S4 for further details). Although these enthalpy estimates are notoriously characterized by large statistical uncertainties (and in spite of the compensatory contribution expected from the entropic term), the Lennard-Jones energy difference is relatively large and suggestive of a significant role of steric interactions in ion selectivity. We thus surmise that the free energy difference between the two states can be explained by the increased size of the solvated ion in the environment of a quasi-rigid pocket.

This result is consistent with the experimental finding that the E2P ion binding site can accommodate acetamidinium, while the E1P ion binding site does not, mostly likely due to size and hydration structure of this organic cation (25). Furthermore, a study by Jorgensen et al. (26) shows that sodium-ion-dependent enzyme phosphorylation is reduced to a greater degree with mutations that change side chain volume but preserve total charge of the side chain moiety, i.e. mutations E327D, E779D and D804E, compared to mutations that change total negative charge but preserve side chain volume, i.e. mutations E327Q, E779Q, and D804N (Table S5). This suggests that the E1P state is sensitive to side chain volume changes, supporting a mechanism where binding site volume controls selectively.

Unlike the E1P state, the higher affinity of the E2P state for potassium ions compared to sodium ions appears to be more dependent on the specific protonation state. In this case, the looser steric restrictions allow either two sodium or potassium ions to bind; however, the clear trend is that, as pH decreases, the relative selectivity for potassium ions increases (Figure 2D). This prediction is consistent with well-established experimental observations (19,20). Hence, we surmise that the protonation state is a key factor in determining the selectivity of the E2P conformation. Next we examine how the E2P state protonation state may be coupled to its conformational state.

#### A cytoplasmic binding site for anions controls the protonation state for D926

To understand how protonation state is controlled in the Na+/K+-ATPase, we used electrostatic potential decomposition to investigate how the protein and/or its environment affects pKa of the five acidic binding site residues (see Experimental procedures). Briefly, for the most probable protonation state in E1P and E2P, we calculated the electrostatic potential of placing a negative charge density on the region occupied by sidechains of residues D804, D808, D926, E327 and E779 (see Experimental procedures). Therefore, in this analysis, relatively positive electrostatic potentials favor deprotonation while relatively negative electrostatic potentials favor protonation (Figure 3A). Comparing the electrostatic potential generated by the entire system with the potential generated only by the protein reveals that the protein follows the same protonation/deprotonation patterns as the ones predicted by the entire system for all binding site residues in both E1P and E2P states, except D926 in the E2P state (Figure 3A, blue box). For D926 in the E2P state, the protein favors deprotonation of this residue, while the system as a whole, favors protonation (relative to the other residues, see Experimental procedures). Therefore, protein conformational change is not affecting electrostatic environment of D926. D926 is of particular importance in function of Na+/K+-ATPase; as we showed earlier, D926 needs to be protonated in the E2P state to facilitate proper potassium binding to sites I and II, and deprotonated in the E1P to allow sodium ion to bind site III. While the protein conformational changes in E1P and E2P states seem to be responsible for protonation/deprotonation of most binding site residues, environmental factors seem to be controlling protonation state of D926. Indeed, by decomposing the electrostatic potential into contributions arising from protein, water, lipid, cations (K^+^ for E2P), and anions (Cl^-^), we found that the electrostatic interactions favoring protonation of D926 in E2P state are largely due to the influence of anions (Figure 3B—Tables S11 and S12).

**FIGURE 3.**
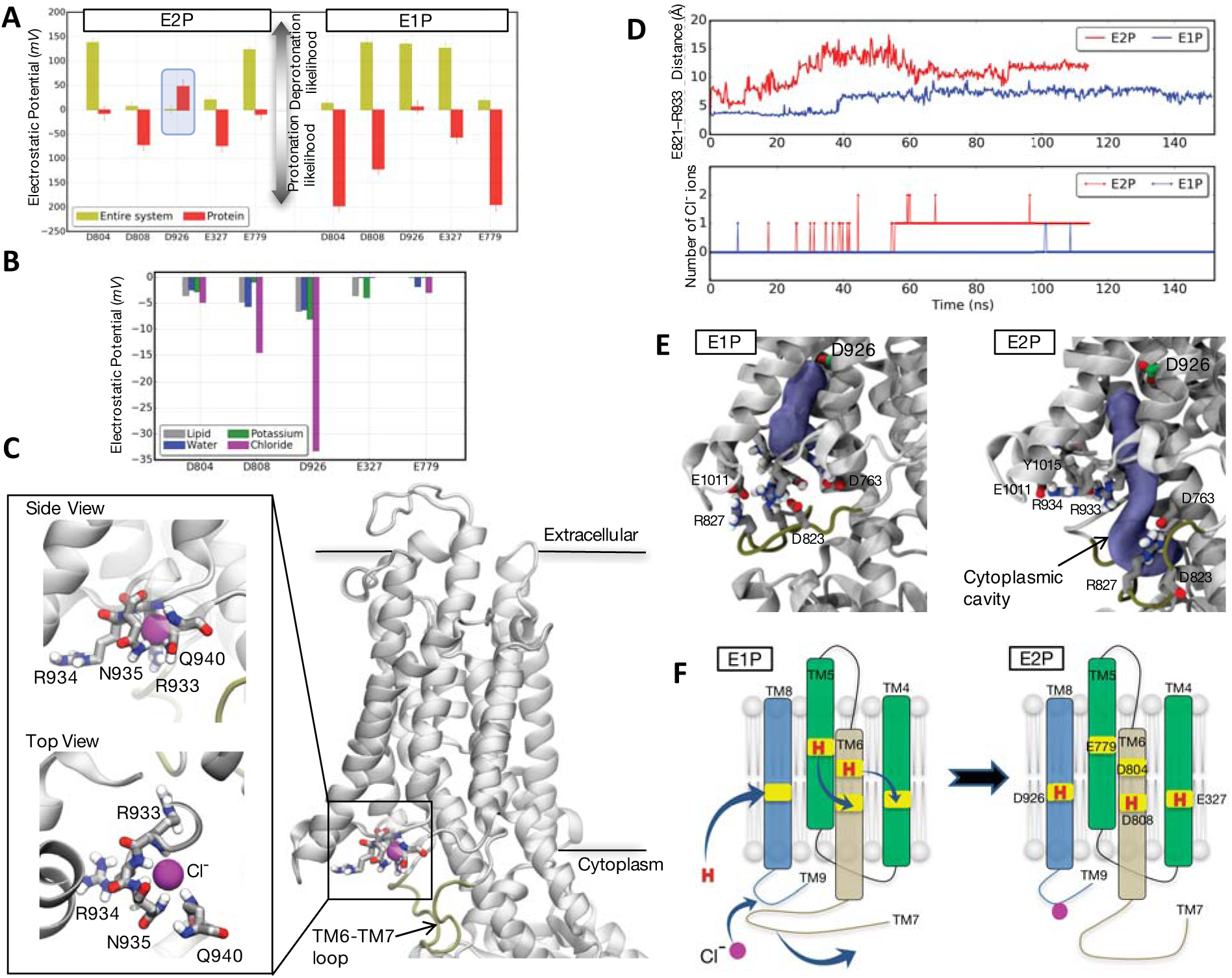
Identification of a putative cytoplasmic anion binding site for Na+/K+-ATPase. (A) Average electrostatic potential due to the protonation of key residues in the Na^+^/K^+^ binding site for both the E1P and E2P states. For each binding site residue, the bars show the electrostatic potential calculated using the entire system (yellow), and the subset of protein atoms (red), without considering contributions from that specific residue (see Experimental procedures). Error bars denote standard deviations. (B) Comparison of the relative contributions of lipids, water, solvent cations (K^+^) and anions (Cl^-^) to the electrostatic potential of key binding residues in the E2P conformation. In each case, the maximum electrostatic potential values are subtracted to properly compare the relative contributions of each component (see Table S12 for values before substraction) (C) Molecular representation of the cytoplasmic anion binding site in Na+/K+-ATPase. Only the α subunit is shown for clarity. The bound chloride ion is shown as a magenta sphere. (D) The distance between the TM6-TM7 loop and the TM8-TM9 loop (top panel) and chloride ion density near the cytoplasmic anion binding site (lower panel) is compared between the E1P and E2P states. The distance is calculated using the C_δ_ atom of E821 and backbone nitrogen of R933. The chloride density is calculated by counting the number of chloride ions within 7 Å of backbone nitrogen of R933. (E) A cavity (blue-gray) extends from D926 (shown in green licorice) to the cytoplasm in the E2P conformation (right), but is blocked in the E1P state (left). The program *fpocket* (51) was used for cavity calculations (see Experimental procedures). (F) Cartoon representation of a proposed mechanism of TM6-TM7 lid opening, leading to the binding of a chloride ion, protonation of D926, and protonation rearrangement of other binding residues.

To understand how anions influence the pKa of D926, the anion density was compared between the E1P and E2P conformations. Calculation of a density difference map near this ion binding residue (Figure S7) reveals a putative cytoplasmic binding site for anions, composed of the side chains of residues N935, Q940, and three backbone NH groups from residues R933, R934 and N935 in the loop between TM8 and TM9 (Figure 3C). In our simulations of the E2P conformation, these residues coordinating a chloride ion on the cytoplasmic face of the enzyme directly below D926 (Figures 3C). Intriguingly, we also found that this binding site has a similar structure to that of crystallographically determined chloride binding sites in other proteins (27,28) (Figure S8). Moreover, our simulations show that in the E1P conformation, access to the anion binding site is blocked by salt bridges formed between arginine, aspartate, and glutamate residues in the TM6-TM7 loop and residues in the TM8-TM9 loop in conjunction with E1011, Y1015 and Y1016 from the C-terminus (Figure S9). These observations suggest that the TM6-TM7 loop acts as a *lid* that gates the chloride binding site, allowing access to anions in the E2P conformation, and obscuring access in the E1P conformation, thus controlling the protonation state of D926 and potentially determining the selectivity of the Na+/K+-ATPase.

Similar to the E1P state, the crystal structure of the potassium-bound E2P state shows close proximity of the TM6-TM7 loop and the TM8-TM9 loop (Figure S10), with E821 from the TM6-TM7 loop interacting with backbone nitrogen atoms of residues R933, R934, and N935; hence, the crystal structures don’t reveal the anion binding site. However, simulations predict dissociation of this flexible loop, the TM6-TM7 loop, (Figure 3C), and association of the TM8-TM9 loop with a chloride ion soon thereafter (Movie 1) only in the E2P state (Figure 3D). We also noticed that after disassociation of TM6-TM7 loop, the site III becomes hydrated via a cytoplasmic cavity that extends from cytoplasm up to residue D926 (Figure 3E). On the other hand, for the E1P state, this cytoplasmic cavity is truncated, hence D926 is not water accessible from the cytoplasm. To confirm that the TM6-TM7 loop disassociation and anion binding is not a singular event and is statistically significant, we initiated 5 independent simulations of the potassium-bound E2P state started from the crystal structure conformation and continued for over 200 ns each (Figure S11). The results show that TM6-TM7 loop always dissociates from TM8-TM9 loop, although it is possible that they re-associate. However, the overall dissociated-to-associated ratio is 4.6 to 1 (Figure S11). Furthermore, simulations show that the chloride ion density increases near the cytoplasmic anion binding site (Figure S11) whenever TM6-TM7 loop is disassociated from TM8-TM9 loop.

In order to directly compare the sodium-bound E1P state with the potassium-bound E2P state, we were limited to use crystal structures obtained from *Sus scrofa*, since the only crystal structures available for the sodium-bound E1P state are from this organism (12,13). Nevertheless, to further investigate the presence of anion binding site for the E2P state, we additionally performed simulations of the high-resolution (2.4 Å) crystal structure of K^+^-bound E2P (PDB code: 2ZXE) from *Squalus acanthias* using our FEP-predicted protonation state for the *Sus scrofa* E2P conformation (residues E334, D815, and D933 in 2ZXE crystal structure are protonated, which correspond to residues E327, D808, and D926 in 3KDP crystal structure, respectively). Similar to the low-resolution (3.5 Å) structure (PDB code: 3KDP), the high-resolution crystal structure of K^+-^bound E2P (PDB code: 2ZXE) shows a glutamate residue side chain (E828 in 2ZXE structure which corresponds to E821 in 3KDP) from the TM6-TM7 loop occupying the anion site (Figure S12). Simulations started from this conformation also showed dissociation of this flexible loop (Figure S13), although the dissociation time (~400 ns) is slower compared to the low-resolution 3KDP structure. Furthermore, the chloride density increases soon after TM6-TM7 loop is dissociated from the TM8-TM9 loop (Figure S13), supporting the existence of the cytoplasmic anion binding site.

For completeness, we also compared simulated ion binding poses for a low-resolution (4.3 Å) structure of the E1P state (PDB code: 4HQJ) and a high-resolution (2.8 Å) structure (PDB code: 3WGU) (see Figure S14). Similar coordination arrangements were found in both sets of simulations.

Poulsen et al. (5) have previously suggested that a cavity between TM5, TM7 and TM8 provides cytoplasmic access in the E2P conformation for protonation of D926 and proton leak currents (29,30). Cytoplasmic access to this cavity was also postulated to be blocked in the E1P conformation by the C-terminal hydrogen-bond network (5). However, their simulations did not provide insight into why accessibility of this cavity could change the pKa of D926. While not negating the importance of hydrogen-bonding in controlling cytoplasmic accessibility of this cavity, our simulations suggest that anion site occupancy is the major factor determining the change in local electrostatic environment that controls the D926 pKa change between the E1P and E2P conformations.

### Inherited mutations and Hofmeister effects support the hypothesis of a cytoplasmic anion binding site

The anion binding event observed in our simulations and its energetic characterization provide a viable explanation for the conformation-dependent pattern of protonated residues. As the system relaxes from the E1P state to the E2P state, the TM6-TM7 lid opens up and a chloride ion binds to the cytoplasmic anion binding site (Figure 4F). The consequences of this binding are twofold: (i) the electrostatic potential in the region of space occupied by D926 is modified in such a way as to favor the protonated form of this residue; (ii) the motion of the lid results in the formation of an aqueous cavity joining D926 with the cytoplasm (Figure 3F).

One of the most immediate predictions of this model is that the structural integrity and/or the sequence identity of the TM6-TM7 and TM8-TM9 loops should be required for proper function of the pump. Intriguingly, inherited mutations of the *ATPA2* gene associated with hemiplegic migraine have been mapped to the TM8-TM9 loop region (31,32). Indeed, a single-residue polymorphism R933P identified in these studies has been shown to dramatically disable the activity of the Na+/K+-ATPase (5,33,34). The mutation R933A has significant, but less severe effects on activity (35), consistent with the loss of the backbone NH in R933P but not R933A. To further probe the effects of this mutation on anion binding, we performed a ~70 ns simulation of E2P R933P starting from a chloride-bound conformation. The chloride ion leaves the binding site within 3 ns, and remains dissociated (Movie 2). These observations do not exclude other explanations for the functional importance of TM8-TM9 loop residues, in particular, the prevailing model which posits that TM6-TM7 loop mutations and C-terminal tyrosine mutations disrupt a hydrogen bonding network critical to the C-terminal cytoplasmic ion pathway (5). Our findings are remarkably consistent with this model, but with the slight difference in interpretation that C-terminal tyrosine interactions with the R933 side chain are crucial in anchoring the putative cytoplasmic anion binding site.

Additional data consistent with the involvement of anions in the function of the Na+/K+-ATPase comes from extensive studies characterizing a Hofmeister effect whereby chaotropic ions stabilize the E1P state (24,36) and slow the E1P-E2P interconversion rate (37). While previous studies have ascribed the Hofmeister effect in the Na+/K+-ATPase and the sarcoplasmic reticulum Ca^2+^-ATPase to modified electrostatic interactions at the membrane/enzyme/cytoplasm interface (37-39), the existence of an anion binding site near this interface is not inconsistent with these data. Future simulation work will help to test this hypothesis by predicting binding affinities for different anionic chemical species.

We also note that the presence of glutamate E821 bound to this site in the crystal structure of the E2P state is not inconsistent with the existence of an anion binding site. Crystallographic conformations are often biased toward low-energy and/or low-entropy configurations. E821 is in a region that is disordered in the crystal and likely highly mobile in the physiological state.

While this manuscript was under review, a study by Rui et. al. (40) was published showing that protonation states for E1P and E2P are different in the binding aspartate residues. They found that D804 is protonated in E1P state but D808 and D926 are protonated in the E2P state, which agrees with our findings. Unlike our study, the authors assumed a protonated state for all glutamate residues without performing FEP calculations. Our FEP calculations determined protonation states for glutamate residues and showed that E779 is protonated in E1P state but E327 is protonated in the E2P state. Furthermore, while Rui et al. attributed ion selectivity to protonation differences, our findings suggest that only the E2P state shows protonation-dependent selectivity, while binding site volume constraints are responsible for controlling selectivity for the E1P state. We also demonstrated that the protonation of D926 is a crucial step in determining ion binding site selectivity and showed that dissociation of TM6-MT7 loop from TM8-TM9 loop creates a binding site for anions that is responsible for pKa shifts of D926 between E1P and E2P states. Intriguingly, another recent study (41) has found that conformational changes in the intracellular domains (see Figure 1) are particularly sensitive to the protonation state of the binding site. This finding allows us to speculate that the TM6-TM7 loop dissociation is coupled to conformational changes in the intracellular domains, which are in turn triggered by ATP binding and hydrolyses.

### Conclusion

The mechanism by which ion selectivity is achieved in Na+/K+-ATPase is a long-standing question. The structural similarity of the sodium/potassium ion binding site in both sodium- and potassium-occluded conformations suggests that the pump utilizes subtle mechanisms for controlling selectivity. Here, we have conducted a large-scale simulation study of Na+/K+-ATPase in both the E1P and E2P conformations to determine that the protonation state is indeed different for sodium-bound versus potassium-bound states. Our modeling suggests the potassium-bound E2P conformation is particularly sensitive to the protonation state, while for the sodium-bound E1P conformation, van der Waals interactions at the binding sites are more important. The change in protonation state for D926 is particularly crucial for proper binding of sodium and potassium ions. Close inspection of our simulation results reveals a putative, previously unknown cytoplasmic anion-binding site whose occupancy promotes the D926 protonation by changing local electrostatic potential and increasing cytoplasmic accessibility of a cavity in the protein that extends to D926. Access to the cytoplasmic anion-binding site is itself controlled by the TM6-TM7 loop, whose disposition is different in E1P vs E2P states, and likely coupled to larger domain motions in the catalytic cycle of the pump.

We emphasize that the model outlined above provides specific predictions testable by further joint experimental and simulation studies. One key prediction is that mutation or truncation of residues in the TM8-TM9 loop should significantly disturb the pump’s activity via changes in orientation and/or affinity of the putative cytoplasmic anion binding site. Another is that the effects of specific Hofmeister ions should correlate with affinities to the cytoplasmic anion binding site. Future studies will be able to directly test these predictions, as well as to probe in detail the requisite conformational motions involved in transitions between E1 and E2 states of Na+/K+-ATPase.

## Experimental procedures

### System preparation

Molecular structures for the sodium- and potassium-bound Na+/K+-ATPase were obtained from the Protein Data Bank (PDB); the PDB codes are: 4HQJ (E1P.ADP state) and 3KDP (E2P state). These structures were chosen because they share the same sequence and are obtained from the same organism: α_1_ and β_1_ isoforms of pig kidney. The CHARMM-GUI web interface (42) was used to insert the protein into a 1-palmytoyl-2-oleoyl-sn-glycero-3-phosphatidylcholine (POPC) lipid bilayer and surrounded by TIP3P water molecules, Cl^-^ and Na^+^ or K^+^ ions for the E1P or E2P state, respectively. We introduced a phosphorylated aspartic acid at position 369 by using parameters present in the CHARMM27 force field for aspartate and phosphate moieties. The total number of particles is 233660 and 261581 for the E1P and E2P systems, respectively. A summary of system specifications is shown in Table 1.

**Table 1.** Summary of the system specifics for molecular dynamics simulations.

|  | Total number<br>of atoms | Number of ions | Number of lipid<br>molecules | Number of water<br>molecules |
| --- | --- | --- | --- | --- |
| Na<sup>+</sup>-E1P | 233660 | Na<sup>+</sup> : 185 , Cl<sup>-</sup> : 164 | 314 | 56953 |
| K<sup>+</sup>-E2P | 261581 | K<sup>+</sup> : 207 , Cl<sup>-</sup> : 185 | 336 | 65179 |

### Molecular dynamics equilibration simulations

The CHARMM27 force field was used for protein and water; CHARMM36 was used for lipids. Periodic boundary conditions were employed for all of the MD simulations and the electrostatic potential was evaluated using the particle-mesh Ewald method (43) with grid spacing of 1.2 Å and non-bonded cutoff of 11 Å. The lengths of all bonds containing hydrogen were constrained with the SHAKE/RATTLE algorithm (44). The system was maintained at a temperature of 300 K and pressure of 1 atm using a Langevin thermostat and barostat, as implemented in the MD code NAMD2.10 (45). The rRESPA multiple time step method was employed (46), with a high frequency timestep of 2.0 fs and a low frequency time step of 4.0 fs. Molecular configurations were saved every 20 ps. Harmonic restraints with a force constant of 5 kcal/mol/Å^2^ were applied on the phosphorus and nitrogen atoms of lipids and protein backbones and released gradually during the first 10 ns of simulations. Simulation times for equilibration runs are shown in tables S2 and S3..

### Metadynamics simulations

Initially, the equilibrated structures in which none of the acidic resides in the binding site were protonated was used for metadynamics simulations to study binding site preferences of Na^+^ and K^+^ ions. Then, D926 was protonated in the E2P state to compare the results with the deprotonated D926 case (see Figure 1 and the main text). In all cases, two *distanceZ* collective variables (22) were used to build the potential of mean force (PMF) and to confine one Na^+^ or K^+^ ion in a rectangular box of 30 × 20 Å^2^ inside the binding site. Potential hills with a height of 0.05 kcal/mol and a width of 0.25 Å were added every 200 steps for a total simulation time of 70 ns.

### FEP calculations to identify the lowest-energy protonation state

Free energy perturbation (FEP) calculations were performed by linearly interpolating the Hamiltonians of the protonated and deprotonated systems: (1-λ)*U*_prot_ + λ*U*_deprot_, with the progress variable λ varying between 0 and 1. For each transformation, trajectories for different values of λ (windows) – each window was simulated for 10-60 ns, were collected. Additional harmonic potentials were added to the potential energy function to keep the particles of the decoupling and coupling amino acids in close proximity. These additional interactions forced contiguous windows to sample overlapping regions of the conformational space, thereby increasing the rates of convergence of the free energy estimates. Subsequent analysis of trajectories was limited to the stable part of each window, i.e. the one with no detectable drift in the potential energy. Simulation times for all FEP runs are shown in Table S6. Initially, 20 window FEPs were used, each window followed from the equilibrated structure of the previous window to increase the rate of convergence. Then, in selected cases (when the ∆∆*G* was affected by large uncertainties), we used 40 FEP windows – all started with the same equilibrated structure obtained from at least 100 ns of unbiased equilibration MD, to estimate the free energy. In all cases, the results of 40 window FEPs were more accurate than 20 window FEPs and reassuringly, they were always in qualitative agreement with those obtained with 20 window FEPs (Tables S7 and S8). Therefore, we performed 40 window FEPs for all transformations and reported the final results presented in Figure 2 based on these FEPs. In both E1P and E2P state, all alchemical transformations except two entailed the transfer of a proton from one binding site residue to another. In these cases, the overall chemical composition of the system is unchanged during the transformation and the estimated free energy difference is directly comparable to the experiments. In the remaining cases, where the total charge of the system changes during FEP, an auxiliary thermodynamic cycle (reference reactions, described below) is considered in which the FEP of a proton dissociation reaction from a three residue peptide is calculated in solution. Then, this FEP is subtracted from the FEP of protein to compensate charge changing effects (see below for more details).

### FEP calculations for reference (solution) reactions

The reference proton dissociation reactions for Glu and Asp amino acids in solution were characterized by using a tri-peptide, Ile-Asp-Leu for Asp protonation and Pro-Glu-Ile for Glu protonation. These sequences were chosen on the basis of the neighboring residues of D804 and E779 in the Na+/K+-ATPase. Both systems were solvated with TIP3P water molecules and minimized before FEP runs. Similar to other FEP calculations, harmonic restrains were applied to keep the particles of decoupling and coupling amino acids in close proximity. 40 windows were used each with 10 ns equilibration time. All other parameters are the same as described above. See Table S9 for the FEP values.

Instead of using a reference reaction to cancel charge changing effects during FEP calculations, one may use protonation/deprotonation of a side chain in the protein that is far from binding site, therefore the charge of the system will stay the same during the FEP. We tested the accuracy of our reference reaction procedure with this alternative approach for the two FEP reactions in the potassium-bound E2P state. We chose D269 from the beta subunit of the Na+/K+-ATPase to protonate/deprotonate during FEP reactions to cancel charge changing effects in simulations that the charge of the binding site changes. D269 from beta subunit (hereafter denoted as D269_B) is far from binding site and in the extracellular side of the membrane, therefore, the effect of this residue on the binding site is most likely miniscule. The results of these FEP reactions are presented in Table S10. In the case of E327-D269_B 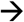 E327-D808 transformation, the FEP result from this alternative approach agrees qualitatively with our reference reaction FEP approach. In the other case, the E327-E779-D269_B 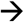 E327-E779-D808 transformation, the result is ambiguous since the error bar for this FEP reaction is more than the FEP value, therefore, the comparison of two approaches couldn’t be achieved. Nevertheless, results of both FEP methods may be interpreted as that the E327-E779 and E327-E779-D808 protonation states have similar free energies and both have higher energy than E327-D808. Therefore, these won’t change the predicted lowest-free-energy protonated state for the E2P conformation shown in Figure 2 of the main text.

### FEP calculations to determine Na^+^ vs K^+^ selectivity

To calculate the free energy difference between the sodium-bound and potassium-bound states, the Lennard-Jones parameters of the bound ion were altered so as to produce a smooth transformation from Na^+^ to K^+^ (Figure S6). Each window was equilibrated for 5 to 10 ns, with the free energy difference between adjacent windows estimated according to the Zwanzig relation ∆*G*(*A→B*) *= -k*_B_*T* ln <exp (-(*E*_*B*_-*E*_*A*_)/*k*_B_*T*)>_*A*_ (47). To ensure its accuracy, we tested this protocol to calculate difference in solvation free energy between sodium and potassium. The result (-19.8 ± 0.1 kcal/mol) is within 1 kcal/mol of previously reported values (-20.7 ± 0.1 kcal/mol) and closer to the experimental value (-17.5 kcal/mol (48)). These FEP values are multiplied by 2 to account for changing 2 potassium ions to two sodium ions in the E2P state and multiplied by 3 to account for changing 3 sodium ions to three potassium ions in the E1P state.

### Analysis of FEP simulations

The Multi-state Bennett Acceptance Ratio (MBAR) method was used for free energy estimation, as implemented in the *pymbar* python package (49,50). Since our FEP calculations to determine the preferred protonation state show good energy overlap not only between adjacent, but also between more distant windows, we used the MBAR estimator, which integrates samples from all thermodynamic intermediates. This estimator is expected to produce superior estimates compared to exponential averaging and/or the (two-state) Bennett Acceptance Ratio (BAR), which only consider adjacent windows.

### Electrostatic potential calculations

To gauge the relative propensity for the deprotonated state of residues D926, E327, E779, D804, and D808, the electrostatic potential energy of a negative charge density located in the regions occupied by each of these residues were estimated. This quantity can be interpreted as the electrostatic component of the free energy difference on changing from protonated to deprotonated state. We estimated this energy using the following equation:

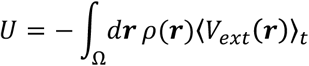

where *ρ*(***r***) is the density of the atoms belonging to side chains of residues D926, E327, E779, D804, or D808; 〈*V*_*ext*_ (***r***)〉_*t*_ is the electrostatic potential, averaged over the entire trajectory, generated by all the charges in the system except those contributing to *ρ*(***r***). Since the system is periodic, the long-range part of the potential has been evaluated using the particle-mesh Ewald method (43) with grid spacing of 1.0 Å and non-bonded cutoff of 11 Å. In this approach, in which the Poisson equation is solved in the reciprocal space, the *k=*0 term is customarily neglected as it is divergent. Therefore, 〈*V*_*ext*_ (***r***)〉_*t*_ is defined only up to an additive constant; we use this property in tables S11 and S12 to define a convenient baseline for the interaction energies. *ρ*(***r***) is estimated from a three-dimensional histogram of atom positions, using Gaussian kernels of width σ_i_ equal to the Van der Waals radius of each particle:

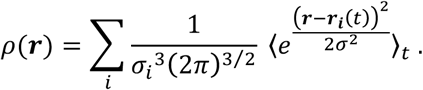

In this equation ***r*** indicates the position in space, ***r***_*i*_(*t*) is the location of the *i*^*th*^ side chain atom at time *t*, the sum runs over all the atoms of the binding site residue under consideration and 〈…〉_*t*_ indicates the average over the MD trajectory. By virtue of linearity of Poisson equation, the external potential (and thus the electrostatic potential) can be decomposed in contributions arising from five distinct, non-overlapping sets of atoms: protein, water, lipid, solution (K^+^ or Na^+^) cations and solution Cl^-^ anions. The electrostatic potentials for each component and for all binding site residues are shown in tables S11 and S12.

### Protein cavity detection

To identify pockets and pathways connecting the binding site to the intracellular milieu we used the program *fpocket* (51,52), which utilizes Voronoi tessellation to define protein volumes. We used the tool MDpocket to analyze the structure of the protein along the trajectory and identified transient and stable pockets (52).

## Acknowledgments

This work was supported in part by the National Science Foundation Grants ACI-1440059 (V.C.), MCB-1412508 (A.R and V.V.) and major research instrumentation grant CNS-09-58854. L.D. receives funding from European Union Seventh Framework Program 979 “Voltsens” Grant PIOF-GA-2012-329534.

## Author contributions

AMR conducted most of the simulations, analyzed data and wrote the initial draft. AMR, VAV, and VC conceived the project and experimental design. All authors participated in interpreting the data and in writing the manuscript.

## The abbreviations used are

MD: molecular dynamics
TM: transmembrane
FEP: free energy perturbation
PMF: potential of mean force
MBAR: Multi-state Bennett Acceptance Ratio
BAR: Bennett Acceptance Ratio

